# Developmental maturation of the hematopoietic system controlled by a Lin28b-*let*-*7*-PRC1 axis

**DOI:** 10.1101/2022.01.05.474696

**Authors:** Dahai Wang, Mayuri Tanaka-Yano, Eleanor Meader, Melissa A. Kinney, Vivian Morris, Edroaldo Lummertz da Rocha, Nan Liu, Stuart H. Orkin, Trista E. North, George Q. Daley, R. Grant Rowe

**Affiliations:** Department of Hematology/Oncology, Boston Children’s Hospital, Boston MA; Stem Cell Program, Boston Children’s Hospital, Boston, MA; Department of Biomedical Engineering, University of Wisconsin-Madison, Madison, WI; Graduate Program of Pharmacology, Center for Biological Sciences, Federal University of Santa Catarina, Florianopolis, SC, 88040-900, Brazil; Howard Hughes Medical Institute, Boston MA; Harvard Medical School, Boston MA; Dana-Farber Boston Children’s Cancer and Blood Disorders Center, Boston, MA; Stem Cell Transplantation Program, Boston Children’s Hospital

## Abstract

Hematopoiesis changes over life to meet the demands of maturation and aging. Here, we find that the definitive hematopoietic stem and progenitor cell (HSPC) compartment is remodeled from gestation into adulthood, a process regulated by the heterochronic Lin28b/*let-7* axis. Native fetal and neonatal HSPCs distribute with a pro-lymphoid/erythroid bias with a shift toward myeloid output in adulthood. By mining transcriptomic data comparing juvenile and adult HSPCs and reconstructing coordinately activated gene regulatory networks, we uncover the Polycomb repressor complex 1 (PRC1) component Cbx2 as an effector of Lin28b/*let-7*’s control of hematopoietic maturation. We find that juvenile *Cbx2*^-/-^ hematopoietic tissues show impairment of B-lymphopoiesis and a precocious adult-like myeloid bias and that Cbx2/PRC1 regulates developmental timing of expression of key hematopoietic transcription factors. These findings define a novel mechanism of epigenetic regulation of HSPC output as a function of age with potential impact on age-biased pediatric and adult blood disorders.

## INTRODUCTION

Normal development and maturation require adaptation of stem cells to shifting physiologic priorities. During prenatal development, hematopoietic stem cells (HSCs) undergo rapid self-renewal to expand and establish the nascent definitive hematopoietic system within the fetal liver (FL) with an erythroid differentiation bias to support growth in the hypoxic uterus (Rebel et al., 1996a; Rebel et al., 1996b; Rowe et al., 2016b). Following birth, juvenile hematopoiesis becomes lymphoid-biased to establish and educate the adaptive immune system (MacKinney, 1978). In later life, hematopoiesis becomes myeloid-biased, an effect that appears to be programmed at the level of HSCs and multipotent progenitors (MPPs)(Pang et al., 2011; Young et al., 2016). These age-related changes in hematopoiesis are paralleled by age biases of blood diseases, including several distinct forms of leukemia and bone marrow failure disorders, illustrating the impact of normal development on disease manifestations (McKinney-Freeman et al., 2012; Rowe et al., 2019).

Recently and historically, much effort has focused on defining the molecular regulators of blood maturation from the fetus to the fully mature adult state. Hematopoietic maturation is associated with dramatic changes in gene expression and chromatin organization in HSPCs (Bunis et al., 2021; Copley et al., 2013; Huang et al., 2016; Rowe et al., 2016b). As an example, the Polycomb repressor complex 2 (PRC2) component Ezh1 is downregulated during the developmental transition from primitive to definitive hematopoiesis to derepress transcriptional programs implementing definitive HSC traits (Vo et al., 2018). As a second example, during maturation of the definitive hematopoietic system, the transcriptional repressor BCL11A controls one of the most well-studied hematopoietic maturation processes - globin switching - by directly silencing human fetal globin (Basak et al., 2020). During definitive hematopoietic development, PRCs are variably required for many aspects of fetal and adult blood formation but the upstream molecular regulation of these heterochronic effects is undefined (Park et al., 2003; Xie et al., 2014).

The heterochronic RNA binding protein Lin28b is a key regulator of definitive hematopoietic maturation. Lin28b has been shown to regulate globin switching, self-renewal of HSCs, and both myeloid and lymphoid maturation via repression of the *let-7* family of microRNAs and modulation of their downstream targets, as well as through *let-7*-indepdendent mechanisms (Basak et al., 2020; Copley et al., 2013; Rowe et al., 2016a; Rowe et al., 2016b; Yuan et al., 2012). Although Lin28b directly and indirectly controls the translation of a multitude of RNAs, the specific mechanisms by which it exerts such profound effects on hematopoiesis have remained elusive. Here, we implicate the *let-7* target and PRC1 component Cbx2 in developmental maturation of HSPCs. We find that HSCs and MPPs redistribute and reprioritize their linage output during development and maturation, and that juvenile lymphoid- and erythroid-biased hematopoietic output requires Cbx2. Cbx2, via regulation of chromatin modifications by PRC1, modulates enhancers controlling master hematopoietic transcription factors for effective developmentally timed expression. These findings reveal how HSPCs mature over time and uncover a novel master axis required for effective hematopoietic maturation.

## RESULTS

### HSPCs undergo developmental maturation

To gain understanding of the developmental maturation of HSPCs, we profiled various populations of primitive hematopoietic stem cells (HSCs) and multipotent progenitors (MPPs) during maturation of the definitive hematopoietic system. We found that within the Lineage^-^ Sca-1^+^ CD117^+^ (LSK) fraction, lymphoid-biased MPP-4 declined progressively over time (Figure 1A-C). In line with the shift away from erythroid toward myeloid output among progenitors during maturation that we reported previously (Rowe et al., 2016b), we found that erythroid-biased MPP-2 diminished while myeloid-biased MPP-3 increased with maturation (Figure 1A-C). We did not observe significant differences in the quantity of phenotypic HSCs save for a short-term (ST)-HSC contraction with aging, consistent with prior reports (Figure 1A-C)(Pang et al., 2011). These changes in HSPC content paralleled the content of the peripheral blood, with neonates possessing higher absolute lymphocyte counts than adults (Figure 1D). We also observed lower red blood cell (RBC) counts and increased mean corpuscular volume (MCV) in neonates relative to adults with an overall lower hemoglobin content (Figure 1D). These findings recapitulate known age-dependent changes in peripheral blood parameters described in humans (MacKinney, 1978; Saarinen and Siimes, 1978; van Gent et al., 2009).

**Figure 1.**
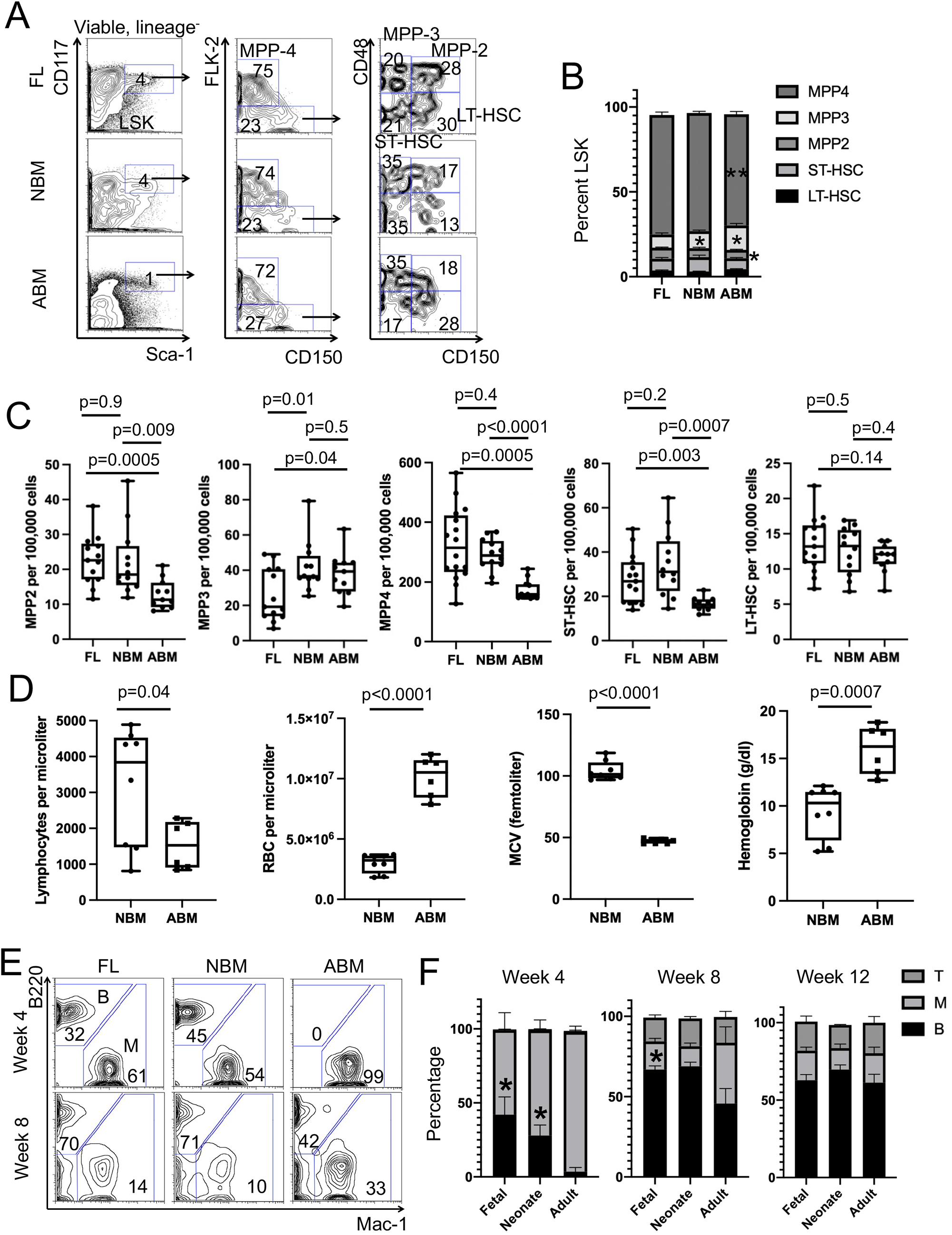
Developmental maturation of HSCs and MPPs. (A) Representative flow cytometry plots demonstrating distribution of HSCs and MPPs in E14.5 fetal liver (FL), neonatal bone marrow (NBM), and adult bone marrow (ABM). (B) Distribution of HSC and MPP populations as a percentage of viable Lineage-Sca-1+ c-kit+ (LSK) populations (* p < 0.05 compared to FL, ** p < 0.05 compared to NBM) (C) Frequencies of various HSC and MPP populations per 100,000 viable cells in each hematopoietic organ. Results aggregated over three independent experimental cohorts. (D) Peripheral blood parameters in NBM and ABM. (E) Representative flow cytometry profiles of the progeny of LT-HSCs from the indicated sources engrafted into congenic recipients at the indicated time points. (F) Quantification of the relative lineage output of LT-HSCs from the indicated ages at the indicated time points following transplantation into congenic recipients. * p < 0.05 compared to adult. Results aggregated over two independent transplantation experiments. Data are presented as mean ± SEM with p values shown.

To functionally validate these lineage priorities ingrained within LSKs, we isolated long-term HSCs (Lineage^-^ CD117^+^ Sca-1^+^ CD127^-^ CD48^-^ CD150^+^; LT-HSCs) from midgestation (embryonic week 14.5 (E14.5)) fetal liver (FL), newborn (postnatal day 0-1 (P0-1)) bone marrow, or young adult (postnatal age 6-8 weeks) bone marrow and transplanted them into congenic adult recipients and monitored lineage output in the donor-derived fraction. We found that E14.5 FL and neonatal LT-HSCs showed significantly more B-cell output at the 4-week time point compared to young adult LT-HSCs, which produced only myeloid cells (Figure 1E-F). After 4 weeks, adult LT-HSCs produced progressively more B-cells, while LT-HSCs of all ages begat T-cells with similar kinetics (Figure 1E-F). Notably, these differences were obscured when we transplanted whole hematopoietic tissue from each of these sources, suggesting that they are programmed at the level of early HSCs (Figure S1).

### HSPC distribution is controlled by the Lin28b/*let-7* axis

Lin28b is a well-established heterochronic factor regulating the timing of developmental events (Shyh-Chang and Daley, 2013). In the hematopoietic system, Lin28b is expressed in the fetal state where it regulates lymphoid differentiation, HSC self-renewal, platelet activation, and myeloid progenitor distributions with its expression decreasing in HSCs with maturation; many of these effects are mediated by repression of the maturation of *let-7* microRNAs(Rowe et al., 2016a). We confirmed that *LIN28B* and several of its known targets decrease with maturation in human and mouse HSPCs (Figure 2A, Figure S2)(Beaudin et al., 2016; Cesana et al., 2018). We also observed that the *LIN28B* promoter is developmentally regulated (Figure S2)(Huang et al., 2016). We next examined the effect of ectopic activation of LIN28B within the adult bone marrow on HSCs and MPPs. Here, we used a double transgenic system for activation of LIN28B in the upon doxycycline treatment for two weeks prior to transplantation (Rowe et al., 2016b). At 12 weeks post transplantation, we found that LIN28B activation tended to diminish MPP-3, increased MPP-4, and diminished LT-HSCs consistent with partial implementation of a fetal HSC/MPP state, suggestive of influence of the adult hematopoietic niche (Figure 2B-C)(Young et al., 2021). Finally, we ectopically expressed a degradation-resistant form of *let-7g* in the adult marrow post transplantation, finding that this altered adult LSKs, with an increase in MPP-3 and diminishment of MPP-2 (Figure 2D-E). Together, these data demonstrate the Lin28b/*let-7* modulates the developmental state of HSCs and MPPs.

**Figure 2.**
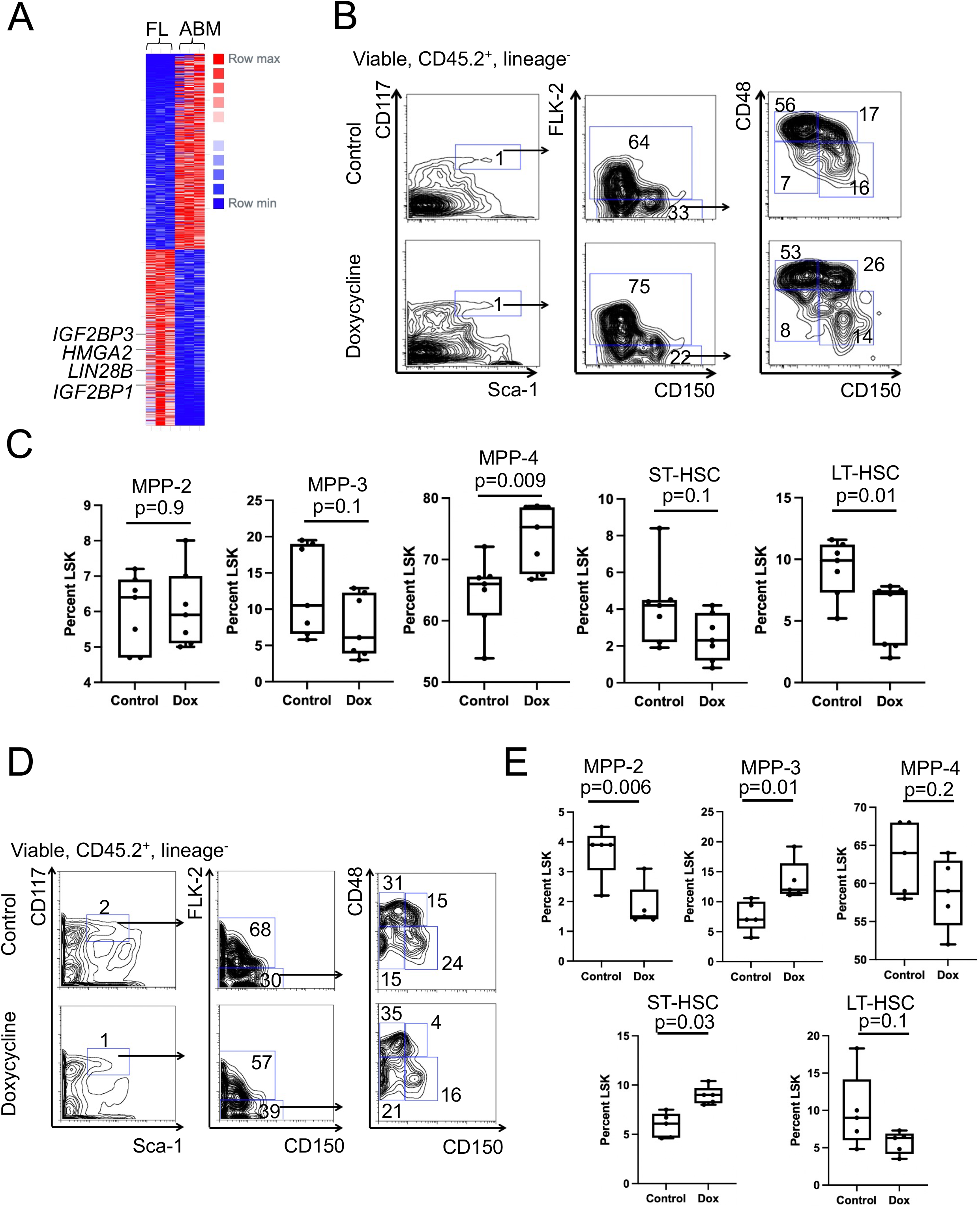
Regulation of HSC and MPP maturation by Lin28b and *let-7* microRNAs. (A) Heatmap demonstrating differential gene expression between FL and adult BM human HSCs (Cesana et al. 2018). (B) Adult BM cells engineered for doxycycline inducible LIN28B expression were transplanted into congenic recipients which were maintained with or without doxycycline in the drinking water. After 8 weeks, the distribution of HSCs and MPPs was examined in the BM by flow cytometry. (C) Quantification of the indicated HSC and MPP populations in control or doxycycline-treated mice at 8 weeks following transplantation. Results are aggregated over two independent experiments. (D) Adult BM cells engineered for doxycycline-inducible, degradation-resistant *let-7g* expression were transplanted into congenic recipients which were maintained with or without doxycycline in the drinking water. After 8 weeks, the distribution of HSCs and MPPs was examined in the BM by flow cytometry. (E) Quantification of the indicated HSC and MPP populations in control or doxycycline-treated mice at 8 weeks following transplantation. In all panels, data are presented as mean ± SEM with p values shown.

### Cbx2 is a developmentally regulated Lin28b/*let-7* target in the hematopoietic system

Next, we endeavored to determine the downstream mediators of Lin28b’s effect on HSPC maturation. We used the CellNet algorithm to query 1787 mouse gene regulatory subnetworks (GRSs) incorporating 717,140 genetic interactions using a published hematopoietic progenitor RNA sequencing (RNA-seq) dataset (Cahan et al., 2014; Rowe et al., 2016b). This dataset includes common myeloid progenitors (CMPs) from E14.5 FL, WT adult bone marrow, and adult bone marrow ectopically expressing LIN28B (iLIN28B) to determine which GRSs are coordinately enriched in E14.5 FL and the FL-like maturation state implemented by ectopic LIN28B (Figure 3A)(Rowe et al., 2016b). We found 23 and 19 GRSs coordinately activated and repressed, respectively, in E14.5 FL and ectopic LIN28B marrow (Figure 3B). Focusing on GRSs active in hematopoietic tissues (blood subnetworks), we found 3 and 5 GRSs activated and repressed, respectively (Figure 3B). We next queried coordinately regulated GRSs for predicted *let-7* microRNA targets using Targetscan, which would be predicted to be increased in LIN28B-expressing cells (Agarwal et al., 2015). Using this approach, we identified the *let-7* target *Cbx2* as enriched in E14.5 FL and iLIN28B adult CMPs compared to *Lin28b*^-/-^ FL CMPs and WT adult CMPs (Figure 3C).

**Figure 3.**
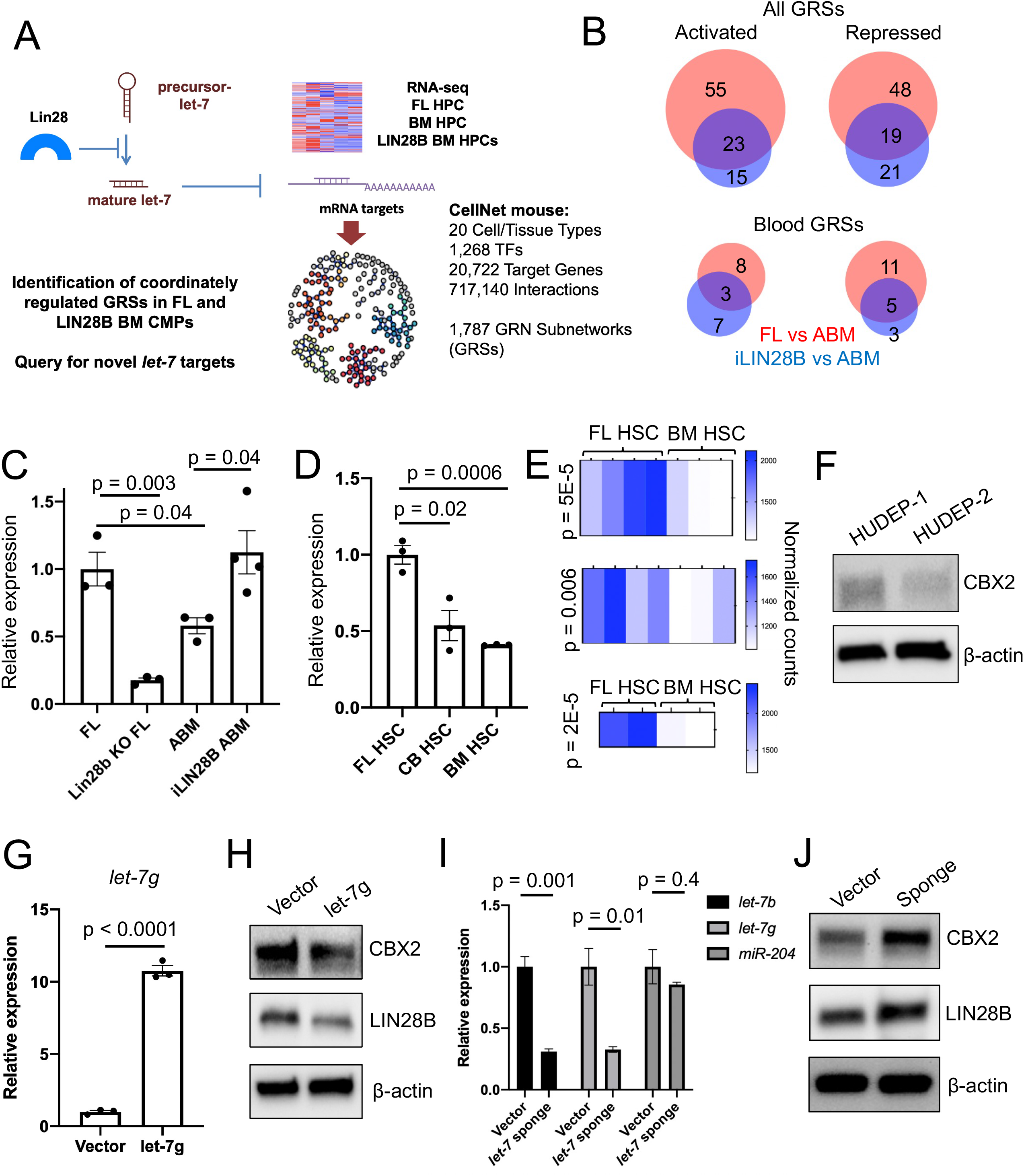
Cbx2 as a candidate Lin28b/*let-7* target. (A) Approach to querying datasets for candidate GRSs that are developmentally regulated by Lin28b and which contain *let-7* target transcripts. (B) Quantification of GRSs coordinately activated or repressed in E14.5 FL compared to ABM progenitors, or LIN28B-expressing ABM progenitors compared to wild-type ABM progenitors (Rowe et al., 2016b). (C) Quantification of *Cbx2* expression in the indicated common myeloid progenitor populations by RNA sequencing (Rowe et al., 2016b). (D) Quantification of *CBX2* expression in the indicated human HSC populations by RNA sequencing (Cesana et al., 2018). In all panels, data are presented as mean ± SEM with p values shown. (E) Heatmaps showing analysis of RNA sequencing data of *Cbx2* expression in two subpopulations of murine FL HSCs compared to adult bone marrow HSCs (top two panels(Beaudin et al., 2016)) or in an independent dataset(Chen et al., 2019; Tober et al., 2018). p values shown. (F) Levels of CBX2 protein were measured by western blotting in HUDEP-1 (neonatal) and HUDEP-2 (adult) erythroid progenitor cells. (G) Expression of *let-7g* was measured in K562 cells expressing the indicated constructs by quantitative RT-PCR. (H) Levels of the indicated proteins were measured in K562 cells expressing the indicated constructs by Western blotting. (I) Levels of the indicated microRNA species were measured by quantitative PCR. (J) Levels of the indicated proteins were measured in K562 cells expressing the indicated constructs by Western blotting. In all panels, results are presented as average ± SEM compared by student’s t-test, with p values shown.

We found that *CBX2* expression was higher in human FL HSCs compared to adult HSCs (Figure 3D)(Cesana et al., 2018). By analyzing datasets comparing gene expression in murine fetal versus adult HSCs, we found that *Cbx2* transcripts were reproducibly higher in fetal cells (Figure 3E)(Beaudin et al., 2016; Chen et al., 2019; Tober et al., 2018). However, we did not observe developmental regulation of the *CBX2* promoter, consistent with its regulation being post-transcriptional (Figure S2)(Huang et al., 2016). Using human erythroid progenitor cells lines derived from neonatal (HUDEP-1) or adult (HUDEP-2) HSPCs, we found that CBX2 protein was higher in HUDEP-1 cells (Figure 3F)(Kurita et al., 2013).

We next sought to determine whether the Lin28b/*let-7* axis could control CBX2 in hematopoietic cells. First, we generated K562 cell lines with stable overexpression of *let-7g* or control cells bearing the empty vector (Figure 3G). We found that overexpression of *let-7g* diminished endogenous CBX2 protein with a concomitant decrease in LIN28B protein -another *let-7* target (Figure 3H). Next, we introduced a *let-7* sponge construct into K562 cells (Kumar et al., 2008), which specifically sequesters mature *let-7* species (Figure 3I). Cells bearing this sponge showed increased CBX2 protein (Figure 3J). Together, these findings indicate that the Lin28b/*let-7* axis controls CBX2 protein levels and suggest that CBX2 (PRC1) are heterochronic regulators of hematopoiesis.

### *Cbx2* regulates juvenile hematopoietic output

*Cbx2* regulates lymphoid proliferation and can perturb HSPCs upon overexpression, but its developmental stage-specific functions have not been investigated (Core et al., 2004; van den Boom et al., 2013). *Cbx2*-null mice develop skeletal anomalies and sex reversal, and typically die perinatally(Core et al., 1997). We therefore generated *Cbx2*^-/-^ embryos and examined definitive FL hematopoiesis. Compared to *Cbx2*^+/+^ and *Cbx2*^+/-^ littermates, we observed significant diminishment of ST-HSCs and a trend toward diminished MPP-3 in *Cbx2*^-/-^ FLs (Figure S3). We did not observe significant differences in MPP-2, MPP-3, or MPP-4 at this time point (Figure S3). However, we observed remodeling of myeloerythroid progenitors away from a fetal erythroid-dominant distribution and toward an adult-like myeloid predominance, consistent with precocious maturation as seen previously in FLs ectopically expressing *let-7*g (Figure 4A-B)(Rowe et al., 2016b). Consistent with this observation, we also observed a decrease in the master erythroid transcription factors *Gata1* and *Klf1* in *Cbx2*^-/-^ neonatal bone marrow and loss of a heme synthesis signature (Figure 4C-D)(Katoh-Fukui et al., 2019).

**Figure 4.**
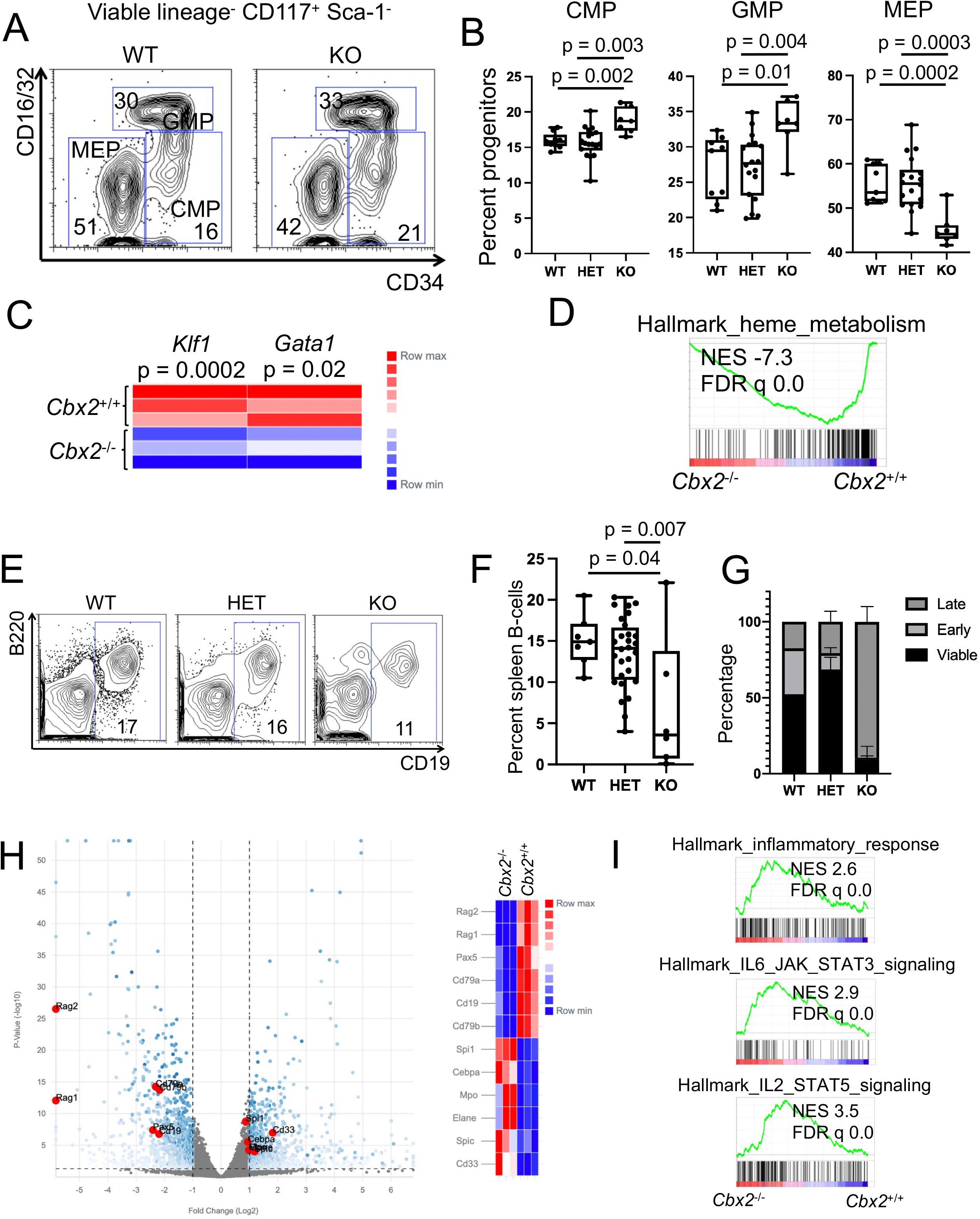
Regulation of juvenile hematopoiesis by Cbx2. (A) Representative flow cytometry plot of viable, lineage-c-kit+ Sca-1-myeloerythroid progenitors from the indicated *Cbx2* genotypes. (B) Quantification of myeloerythroid progenitors from the indicated genotypes. Data are presented as mean ± SEM with p values shown. (C) Heatmap showing expression of erythroid transcription factors in *Cbx2*^+/+^ versus *Cbx2*^-/-^ neonatal bone marrow (Katoh-Fukui et al., 2019). (D) Gene set enrichment analysis of *Cbx2*^+/+^ versus *Cbx2*^-/-^ neonatal bone marrow(Katoh-Fukui et al., 2019). (E) Representative flow cytometry plots of B-cell markers in the P0 neonatal spleen of the indicated genotypes. (F) Quantification of B220+ CD19+ mature B-cells in the P0 neonatal spleen of the indicated genotypes. (G) Quantification of CD19+ B-cells either viable or in the early or late phases of apoptosis. HET viable compared to KO p = 0.0004. (H) Volcano plot and heatmap showing differentially expressed transcripts in *Cbx2*^+/+^ versus *Cbx2*^-/-^ neonatal bone marrow (Katoh-Fukui et al., 2019). (I) Gene set enrichment analysis of *Cbx2*^+/+^ versus *Cbx2*^-/-^ neonatal bone marrow(Katoh-Fukui et al., 2019). In all panels, data are presented as mean ± SEM with p values shown.

We next examined mature blood cell output. We first asked whether *Cbx2* regulates the youthful wave of lymphopoiesis. We found that the *Cbx2*^-/-^ neonatal spleen was deficient in B-cells **(Figure 4E-F**). *Cbx2*^-/-^ B-cells did not undergo precocious maturation from a fetal B-1a to a mature adult B-2 state, but rather showed impaired maturation (Figure S4). B-cells were undergoing apoptosis, likely contributing to the impaired maturation (Figure 4G). RNA sequencing analysis revealed a decrease in transcripts encoding key B-cell factors including *Pax5* in *Cbx2*^-/-^ marrow with a relative gain of myeloid genes - including the master myeloid transcription factor *Spi1*/*Pu*.*1* and signatures associated with inflammatory responses (Figure 4H-I)(Katoh-Fukui et al., 2019).

We next directly examined the functional role of Cbx2 in governing HSPC differentiation. First, we transplanted *Cbx2*^-/-^ MPP-4 isolated from E14.5 FL which resulted in a diminishment of donor-derived B-lymphoid output compared to littermate *Cbx2*^+/+^ FL MPP-4 (Figure 5A-B). Conversely, activation of Cbx2 in adult HSPCs, where it would be otherwise developmentally downregulated, resulted in skewing of cell output toward the erythroid lineage within GEMM colonies, consistent with a fetal-like phenotype **(Figure 5C**)(Rowe et al., 2016b). Finally, we turned to the zebrafish system where we found that morpholino-mediated knock-down of *cbx2* in the developing embryo resulted in diminishment of *rag2:gfp*^+^ lymphocytes in the thymus by 96 hours post fertilization, recapitulating the phenotype observed in mice (Figure 5D-E). Together, these results indicate that *Cbx2* regulates developmental age-specific hematopoiesis.

**Figure 5.**
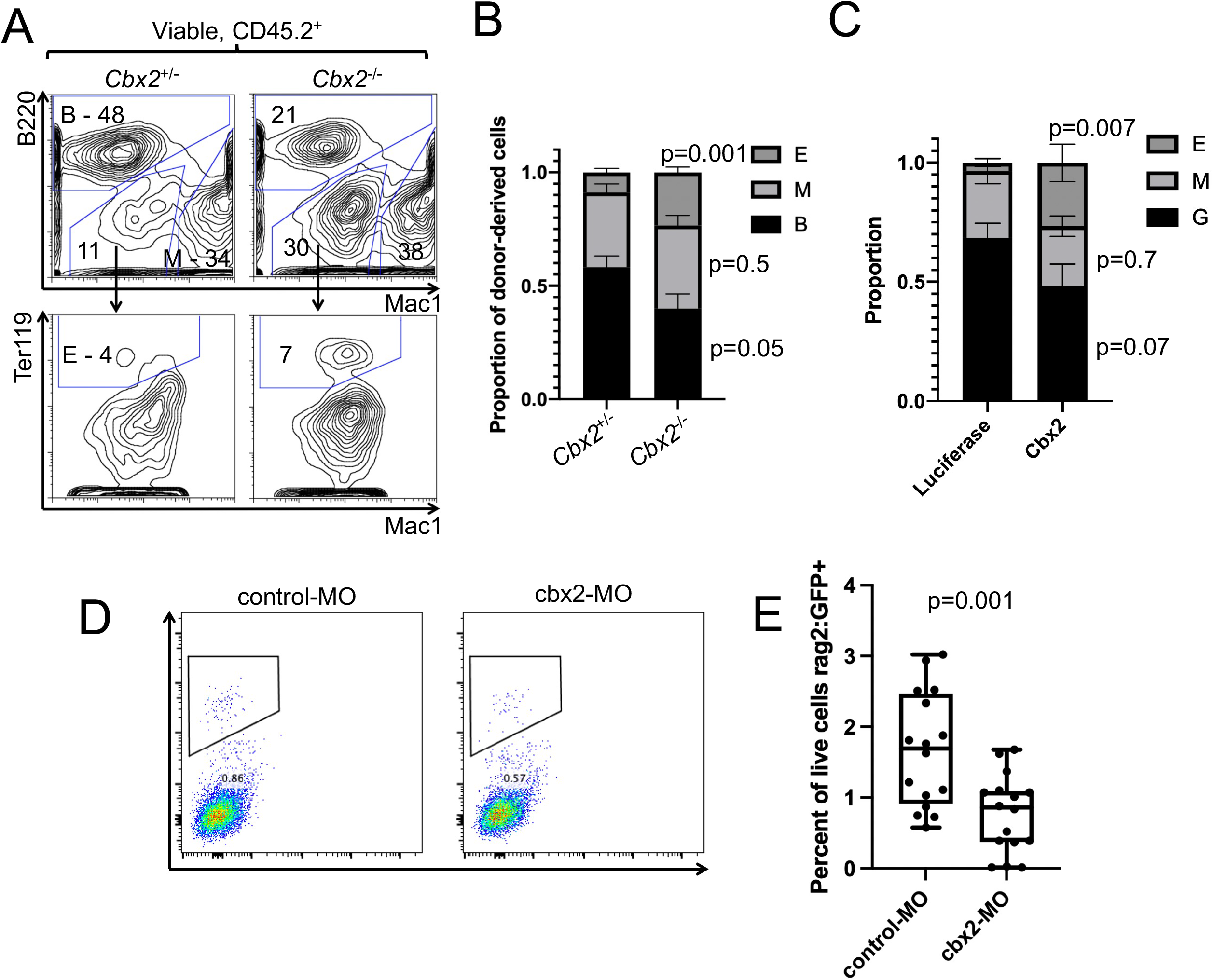
Regulation of lineage output by Cbx2. (A-B) MPP-4 from E14.5 FL were transplanted into congenic recipients. After two weeks, lineage output was analyzed (n = 5 *Cbx2*^+/-^ and *Cbx2*^-/-^ recipients tested. (C) Wild-type adult LSKs were transduced with the indicated vectors and plated in methylcellulose. GEMM colonies were picked and lineage output analyzed by flow cytometry (n = 14 luciferase and 12 Cbx2 colonies). (D-E) Control or *cbx2* targeting morpholinos were introduced into *rag2:GFP* transgenic zebrafish embryos. At 96 hours post fertilization, embryos were dissociated (5 embryos pooled for each data point) and analyzed by flow cytometry. Results are pooled from two independent experiments. In all panels, results are presented as average ± SEM compared by student’s t-test, with p values shown.

### Cbx2 controls PRC1 activity in fetal HSPCs

To understand the role of Cbx2/PRC1 in juvenile hematopoiesis, we performed CUT&RUN for histone H2A lysine 119 monoubiquitinylation (H2AK119Ub), the hallmark of gene silencing by PRC1 required for maintenance of repression of target loci(Skene and Henikoff, 2017; Tamburri et al., 2020). First, we performed H2AK119Ub CUT&RUN using neonatal HUDEP-1 cells. We observed that H2AK119Ub distributed throughout gene bodies relative to histone H3 lysine 4 trimethylation (H3K4me3), which was localized to promoters as expected (Figure S5). We observed H2AK119Ub binding to the pro-myeloid transcription factors *GFI1* and *SPI1*, both of which showed higher levels of the activating histone mark histone H3 lysine 27 acetylation (H3K27Ac) in human adult relative to fetal HSPCs (Figure S5).

To determine the functional role played by Cbx2 in developmental control of PRC1, we next isolated E14.5 FL HSPCs from *Cbx2*^-/-^ embryos or *Cbx2*^+/+^ littermate controls for CUT&RUN analysis. Here, we did not observe global disruption of H2AK119Ub localization in *Cbx2*^-/-^ cells (Figure 6A). We identified several H2AK119Ub peaks gained or lost in *Cbx2*^-/-^ cells relative to controls (Figure 6B). Gene ontology analysis of differential peaks revealed enrichment for several terms related to hematopoiesis, particularly terms related to adaptive immunity (Figure 6C). Motif analysis of peaks diminished in *Cbx2*^-/-^ HSPCs identified binding motifs for hematopoietic transcription factors (Figure 6D).

**Figure 6.**
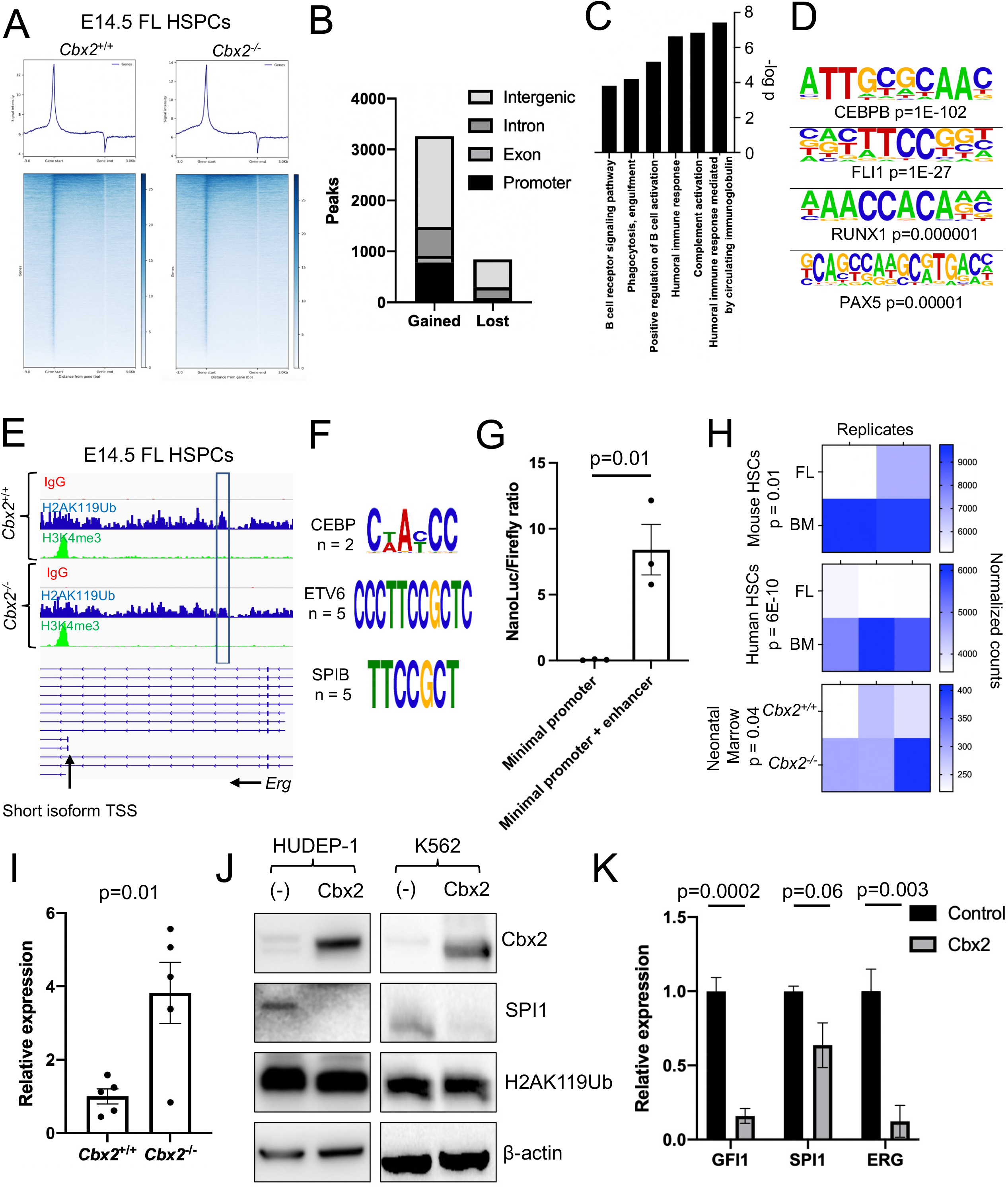
Control of juvenile hematopoiesis by Cbx2/PRC1. (A) CUT&RUN for H2AK119Ub in *Cbx2*^+/+^ compared to *Cbx2*^-/-^ E14.5 FL HSPCs showing distribution in gene bodies. (B) H2AK119Ub CUT&RUN data was compared between *Cbx2*^+/+^ and *Cbx2*^-/-^ E14.5 FL HSPCs to identify differential peaks. (C) Gene ontology analysis of genes associated with peaks gained or lost in *Cbx2*^-/-^ compared to *Cbx2*^+/+^ HSPCs. (D) HOMER analysis of motifs enriched in H2AK119Ub peaks lost in *Cbx2*^-/-^ compared to *Cbx2*^+/+^ HSPCs. (E) CUT&RUN tracks for the indicated markers within the mouse *Erg* gene. Transcriptional start site of the short *Erg* isoform and the candidate enhancer region are indicated. (F) Representative cis binding motifs for the indicated transcription factors identified in the *Erg* enhancer. (G) A fragment of the candidate *Erg* enhancer boxed in (E) was cloned into a vector with a minimal promoter driving expression of Nanoluc. This vector was cotransfected with a constitutive Firefly luciferase vector into K562 cells. 24 hours later, Nanoluc and Firefly signals were collected and expressed as a Nanoluc:Firefly ratio (n = 3 independent experiments). (H) Heatmaps showing *ERG*/*Erg* expression in the indicated populations by RNA sequencing. p-values are shown. (I) Quantitative RT-PCR for *Erg* from neonatal spleens from the indicated genotypes normalized to actin. (J) Western blotting for the indicated proteins in either control HUDEP-1 cells or cells transduced with a Cbx2 expression vector or K562 cells with or without Cbx2 expression from a doxycycline-inducible vector. (K) Quantitative PCR for the indicated transcripts in either control HUDEP-1 cells or cells transduced with a Cbx2 expression vector. In all panels, results are presented as average ± SEM compared by student’s t-test, with p values shown.

We next focused on peaks lost in *Cbx2*^-/-^ HSPCs as potential PRC1 targets directly dependent on Cbx2. Notably, we did not observe differences in H2AK119 monoubiquitylation at the *Hoxa* or *Hoxb* clusters (Figure S5). However, we observed differential H2AK119Ub abundance associated with a 1.4 kilobase (kb) intronic sequence within the *Erg* gene located 52 kb upstream of the transcriptional start site of the short *Erg* isoform (Figure 6E). Analysis of H3K4me3 revealed that the promoter of short isoform was active, consistent with predominant utilization of this isoform in FL HSPCs (Figure 6E). We found several consensus binding sequences for CEBP, ETV6, and SPIB in this within this interval, suggestive of enhancer activity (Figure 6F). We cloned a fragment of this candidate enhancer, finding that it strongly enhanced transcription from a minimal promoter (Figure 6G). Supportive of its developmental regulation by Cbx2, *ERG*/*Erg* is expressed at higher levels in adult BM versus FL human and mouse HSPCs, *Erg* is increased in *Cbx2*^-/-^ compared to *Cbx2*^+/+^ neonatal bone marrow, and the human *ERG* locus showed differential H3K27Ac in adult and FL HSPCs at apparent developmentally regulated candidate human enhancer sequences associated with H2AK119Ub (Chen et al., 2019; Huang et al., 2016; Katoh-Fukui et al., 2019; Tober et al., 2018)(Figure 6H, Figure S5). We confirmed that *Erg* expression is increased in perinatal spleens of *Cbx2*^-/-^ mice compared to *Cbx2*^+/+^ littermates (Figure 6I). Ectopic expression of Cbx2 resulted in repression of SPI1 protein levels and also repressed *SPI1, ERG* and *GFI1* transcripts (Figure 6J-K). Taken together, these data support Cbx2/CBX2-mediated developmental regulation of *Erg*/*ERG* and other master hematopoietic transcription factors. Our results describe a mechanism by which Cbx2, via control of PRC1, regulates hematopoietic maturation (Carmichael et al., 2012; Tsuzuki et al., 2011).

## DISCUSSION

Here, we find that definitive HSPCs undergo stereotypical remodeling during development and maturation from the fetus to the neonate and into young adulthood. We observed that HSCs and MPPs redistribute to reflect age-specific patterns of cell output, from juvenile erythroid- and lymphoid-biased output to mature myeloid predominance, demonstrating that age-associated lineage biases are ingrained early in hematopoietic differentiation. These results extend prior work on postnatal remodeling of the HSC/MPP compartment during aging from young to older adulthood by showing that the aging is preceded by a scripted process of maturation(Young et al., 2016).

The Lin28b/*let-7* axis is the most thoroughly investigated regulator of age-specific hematopoiesis (Copley et al., 2013; Rowe et al., 2016a; Rowe et al., 2016b; Yuan et al., 2012). Lin28 paralogs are highly conserved heterochronic factors that control the schedule of developmental events in several species (Kiontke et al., 2019; Moss et al., 1997). Lin28b exerts wide-ranging effects in HSPCs, with its activity sufficient to impart juvenile hematopoiesis in adult cells either via inhibition of *let-7* microRNA stability or by directly regulating translation of specific mRNAs (Basak et al., 2020; Lee et al., 2013; Rowe et al., 2016b). Lin28b is downregulated during progression of development and maturation, releasing *let-7* microRNAs to implement adult hematopoiesis (Rowe et al., 2016b). As an oncofetal factor, Lin28 proteins have been implicated in the pathobiology of hematologic malignancies, suggesting that Lin28b might recruit fetal-specific HSPC traits in leukemia (Emmrich et al., 2014; Helsmoortel et al., 2016; Manier et al., 2017; Rao et al., 2012).

Here, we find that lymphoid-biased MPP-4 diminish with maturation, and that this process is controlled by Lin28b. Lin28b acts as a heterochronic regulator of lymphopoiesis, where its ectopic expression in mature adult HSCs can induce fetal-like lymphopoiesis characterized by γδ-T-cell, B-1a, and marginal zone B cell production (Yuan et al., 2012). Subsequent studies have shown that the transcription factor Arid3a is expressed in fetal hematopoiesis and is a target of *let-7*; ectopic Arid3a can reprogram adult pro-B cells to produce fetal B-1 cells (Zhou et al., 2015). Lin28b cooperates with Igf2bp3 to stabilize both Pax5 and Arid3a to implement fetal B-lymphopoiesis (Wang et al., 2019). Downstream of differentiation, Lin28b functions in neonatal B-cell positive selection (Vanhee et al., 2019). Our results add to these observations in that we find that not only does Lin28 implement juvenile B-lymphoid differentiation states, but also controls juvenile lymphoid-biased hematopoiesis at the level of HSCs/MPPs, at least in part through Cbx2/PRC1.

Fetal and neonatal hematopoiesis are associated with the production of transient innate-like lymphocyte states, followed by a quantitative burst of lymphoid output presumably to establish innate immunity upon birth. It is well known that the lymphocyte count decreases with aging from childhood to adulthood (Falcao, 1980; MacKinney, 1978). The human thymus forms and is populated by developing lymphocytes undergoing selection in midgestation but begins the process of involution in the first decade of life (Farley et al., 2013; Palmer et al., 2018). It has been hypothesized that dysregulation of this crucial period of postnatal immune education contributes to the risk of developing childhood lymphoblastic leukemia, illustrative of the unique properties of this developmental window (Greaves, 2018). Our work demonstrates that timing of this juvenile lymphopoietic wave is programmed into HSCs/MPPs by the Lin28b/*let-7*/Cbx2 axis.

Through analysis of collections of GRSs cross-referenced with *let-7* targets, we identified Cbx2 as a novel regulator of hematopoietic maturation. Recently, a Lin28a-*let-7*-Cbx2 axis was suggested to control skeletal formation via control of *Hox* gene expression (Sato et al., 2020). Although many *Hox* genes regulate the function of normal and malignant HSPCs, we did not observe developmental regulation of *Hox* loci in HSCs and MPPs (Smith et al., 2011; Yu et al., 2014). We demonstrate alterations of *Cbx2* gene expression with age at the transcript level in HSC/MPPs and at the protein level in HSPCs lines derived from newborn or mature donors in response to modulation of Lin28b/*let-7* (Kurita et al., 2013). In addition to its role in promoting juvenile lymphoid output, the Lin28b/*let-7* axis fine tunes myeloerythroid output for age-appropriate physiology (Rowe et al., 2016b). We find that *Cbx2*^-/-^ mice show a precocious adult-like myeloid-biased progenitor distribution, consistent with its role as an effector of Lin28b’s control over hematopoietic maturation. Our data suggest that this is due at least in part to Cbx2’s ability to modulate the expression key HSPC transcription factors that likely serve to program lineage preferences within HSCs and MPPs. These data define Cbx2 as a heterochronic factor within the hematopoietic system that participates in defining age-specific lineage outputs at the level of HSPCs.

Prior to our study, Cbx2 had been previously implicated in both hematopoiesis and lymphopoiesis. The initial characterization of *Cbx2*^-/-^ mice reported that these mice showed involution of the thymus and small splenic size at 3-4 weeks of age, with impaired proliferation of splenocytes (Core et al., 1997). Altered B- and T-cell differentiation was subsequently reported in *Cbx2*^-/-^ mice (Core et al., 2004). These findings further support a crucial role for Cbx2 in the juvenile lymphoid expansion. Knockdown of *CBX2* in human umbilical cord blood CD34^+^ cells resulted in impaired HSPC self-renewal and skewing away from erythroid and toward myeloid output, paralleling our findings (van den Boom et al., 2013). Relative to lineage-restricted progenitors, Cbx2 is most enriched in HSCs and B- and T-lymphocytes, and its overexpression inhibits clonogenesis and repopulation in transplantation assays, suggesting that its activity must be finely balanced (Klauke et al., 2013). While prior studies have defined roles for Cbx2 in HSC and MPP function, and our work places the hematopoietic functions of Cbx2 within the context of normal development, maturation, and aging and place Cbx2 as a critical downstream regulator of the heterochronic Lin28/*let-7* pathway.

It is becoming increasingly apparent that PRCs play important roles in the timing of hematopoietic development and maturation phenotypes and are also dysregulated in blood malignancies. PRCs likely play key roles in the regulation of stage-specific enhancer usage (Huang et al., 2016). Loss of *Ezh1* in the mouse embryo results in precocious unlocking of hallmarks of definitive HSCs in primitive hematopoietic progenitors (Vo et al., 2018). Deletion of *Ezh2* in FL HSCs causes hematopoietic failure associated with reduction in H3K27me3, while its loss in adult BM HSCs results in a much less severe phenotype, likely due to programmed upregulation of Ezh1, which can compensate for Ezh2 loss (Mochizuki-Kashio et al., 2011). Genetic ablation of *Eed*, which disrupts both Ezh1- and Ezh2-containing PRC2 complexes is tolerated by FL hematopoiesis but depletes adult BM HSCs (Xie et al., 2014). Deficiency of the PRC1 component *Bmi1* is tolerated by FL HSCs at steady state, but the BM of adult mice is progressively depleted of HSCs apparently due to dysregulated self-renewal (Park et al., 2003). Here, we define how the Lin28b/*let-7* axis - a master regulator of hematopoietic maturation - integrates with PRC1 to control age-specific hematopoietic output through epigenetic regulation, explaining how Lin28b exerts such broad effects within the hematopoietic system as a master regulator of HSPC developmental maturation. These findings deepen understanding of how the hematopoietic system changes with age and have potential translational impact to understanding the mechanisms by which blood disorders are biased toward particular ages.

## ACKNOWLEDGEMENTS

This work was supported by the National Institute of Diabetes and Digestive and Kidney Diseases (K08 DK114527-01 to R.G.R.) and the National Heart, Lung, Blood Institute (U01 HL134812 to G.Q.D.).

## AUTHOR CONTRIBUTIONS

Conceptualization, R.G.R.; Methodology, R.G.R.; Formal analysis, R.G.R. N.L., E.L.d.R., M.K.; Investigation, D.W., M.Y., E.M., E.L.d.R. and R.G.R.; Resources, T.E.N.; Data curation, R.G.R, and E.L.d.R.; Writing - original draft, R.G.R.; Writing - revising and editing, R.G.R.; Supervision, T.E.N., G.Q.D., and R.G.R.; Project administration, T.E.N., G.Q.D., and R.G.R.; Funding acquisition, G.Q.D. and R.G.R.

## STAR METHODS

### Resource availability

#### Lead contact

Further information and requests for resources and reagents should be directed to and will be fulfilled by the lead contact, Grant Rowe

#### Materials availability

Plasmids generated in this study will be deposited to Addgene prior to the date of publication.

#### Data availability

Next generation sequencing data (including de-identified human data) will be deposited in GEO and dbGAP with availability at the date of publication.

### Experimental model and subject details

Animals were utilized in accordance with approvals from the Boston Children’s Hospital Institutional Animal Care and Use Committees.

#### Mice and transplantation studies

C57BL/6J (CD45.2) and SJL (CD45.1) mice were from Jackson Laboratory. Timed pregnancies were used to isolate FL cells on postcoital day 14.5 and neonatal day 0-1. 6-8-week-old mice were used for comparison. *Cbx2*^*-*/-^ mice (M33-) were from Jackson Laboratory (stock 006002)(Core et al., 1997). For HSC transplantations, mice were conditioned with 975 rad prior to injection of CD45.2 donor cells via the tail vein into CD45.1 recipients. For progenitor transplants, mice were conditioned with 675 rad prior to injection.

#### Zebrafish use and analysis

Validated splice-blocking morpholino oligonucleotides (GeneTools) targeting *cbx2* [5’-TAGTTTCCTGAGAGAGGAACACAAA-3’] were injected (1-2nl of 50µM MO) at the 1-cell stage as previously detailed (Cortes et al., 2016; Huang et al., 2013). Flow cytometry was performed using transgenic *Tg(rag2:GFP)* embryos at 96 hpf (Traver et al., 2003). Embryos (pools of 5 embryos per sample x 8 replicates) were dissociated and analyzed following staining with SYTOX Red viability stain (ThermoFisher).

#### Cell culture

HUDEP-1 (RRID:CVCL_VI05) and HUDEP-2 (RRID:CVCL_VI06) cells were obtained from RIKEN and maintained in SFEM (Stem Cell Technologies) supplemented with 50 ng/ml SCF, 3 units/ml recombinant erythropoietin, 1 µg/ml doxycycline, and 1 µ*M* dexamethasone. K562 cells (RRID:CVCL_0004) were maintained in IMDM with 10% fetal calf serum.

### Method details

#### Flow cytometry

The following antibodies were used (all from Biolegend): CD3 (clone 17A2), Ter119 (TER-119), Gr-1 (RB6-8C5), B220 (RA3-6B2) conjugated to Pacific Blue, CD117 APC-Cy7 (2B8), Sca-1 PE-Cy7 (D7), Flk2 PE (A2F10), CD48 FITC (HM48-1), CD150 APC (TC15-12F12.2), B220 PE-Cy7 (RA3-6B2), Mac-1 PE-Cy5 (M1/70), CD3 PE (17A2), Ter119 APC-Cy7 (TER-119), CD45.2 FITC (104), CD45.1 APC-Cy7 (A20), CD16/32 PerCP-Cy5.5 (93), CD34 FITC (RAM34), CD5 APC-Cy7

(53-7.3). CD19 PE (eBio1D3) was from eBioscience. Cells were labeled with Sytox Blue viability stain (Thermo). Data were acquired on LSRII or LSR Fortessa instruments (BD Biosciences). APC-annexin V was from Biolegend.

#### Gene regulatory subnetwork analysis

Annotated gene regulatory subnetworks generated by the CellNet algorithm(Cahan et al., 2014) were used in gene set enrichment analysis against preranked lists generated from RNA-sequencing-based comparison of WT fetal liver CMPs, WT young adult CMPs, or young adult CMPs ectopically expressing LIN28B (Rowe et al., 2016b). Subnetworks with positive enrichment scores were examined for predicted *let-7* target sites using the Targetscan algorithm (Agarwal et al., 2015).

#### Western blotting

The following antibodies were used: CBX2 (Abcam 80044; Bethyl A302-524A), LIN28B (Cell Signaling Technologies 11965, clone D4H1), SPI1/PU.1 (Cell Signaling Technologies 2258), H2AK119Ub (Cell Signaling Technologies 8240) and β-actin (Cell Signaling Technologies 4970).

#### Quantitative PCR

Primer assays for microRNA species were purchased from Qiagen. Human *CBX2* primers were: F: 5’-GACAGAACCCGTCAGTGTCC-3’, R: 5’-GGCTTCAGTAATGCCTCAGGT-3’. Human *GAPDH* primers were F: 5’-GTCTCCTCTGACTTCAACAGCG-3’, R: 5’-ACCACCCTGTTGCTGTAGCCAA-3’. Human *GFI1* primers were: F: 5’-GAGCCTGGAGCAGCACAAAG-3’, R: 5’-GTGGATGACCTCTTGAAGCTCTTC-3’. Human *SPI1* primers were: F: 5’-GACACGGATCTATACCAACGCC-3’, R: 5’-CCGTGAAGTTGTTCTCGGCGAA-3’. Human *ERG* primers were: F: 5’-GGACAGACTTCCAAGATGAGCC-3’, R: 5’-CCACACTGCATTCATCAGGAGAG-3’. Mouse beta-actin primers were: F: 5’-ACGAGGCCCAGAGCAAGAGAGG - 3’ and R: 5’ - ACGCACCGATCCACACAGAGTA - 3’. Mouse *Erg* primers were: F: 5’ - GAGTGGGCGGTGAAAGAATA - 3’ and R: 5’ - TCAACGTCATCGGAAGTCAG - 3’.

#### Molecular cloning

pMSCV-neo let-7g was a gift from Tyler Jacks (Addgene plasmid # 14784)(Kumar et al., 2007). pMSCV-puro let-7 sponge was a gift from Phil Sharp (Addgene plasmid # 29766)(Kumar et al., 2008). For ectopic Cbx2 expression, the mouse Cbx2 open reading frame cDNA (Genecopoieia) was cloned into pCW57.1 (gift from David Root; Addgene plasmid # 41393). To generate the *Erg* reporter construct, 162-mer double-stranded oligonucleotides containing Erg enhancer sequences were inserted into the pNL3.1 Vector (#N1031, Promega, USA) to establish the pNL3.1-Erg vector.

#### Reporter assays

K562 cells were co-transfected with pGL4.50 vector (Promega, USA) and pNL3.1 or pNL3.1-Erg. Twenty-four hours after transfection, cells were analyzed for luciferase activity by Nano-Glo Dual-Luciferase Reporter Assay System (Promega, USA). The normalized signal for firefly luciferase activity (NanoLuc luciferase activity/firefly luciferase activity) was normalized to the signal of non-transfected wells.

#### CUT&RUN

CUT&RUN was done according to the manufacturer’s protocol. For mouse HSPCs, raw data were processed using FastQC and CUT&RUNTools and mapped to mm10 (Zhu et al., 2019). Parameters of individual software called by CUT&RUNTools, including Bowtie2, Trimmomatic, Picard (http://broadinstitute.github.io/picard/), Samtools were left unchanged. *E. coli* spike-in DNA was used for normalization. We mapped the sequencing reads to E. coli genome using Bowtie2 (--end-to-end --very-sensitive --no-overlap --no-dovetail --no-mixed --no-discordant) and calculated the “spike_in_ratio” in each sample as “Ecoli_reads/Total_reads”. The bigwig tracks were generated using bamCoverage from deeptools package with “scale_factor” calculated as 1/(spike_in_ratio * cell_number_factor), where “cell_number_factor” is 1 for WT cells and 1.711 for KO cells. For human HUDEP-1 and K562 cells, analysis was performed on the Basepair platform with alignment done with Bowtie2 (basepairtech.com). Differential peak calling was completed with MACS2 and motif analysis was done with HOMER.

### Quantification and statistical analysis

Statistical analysis was performed in Prism and Excel. The statistical details of individual experiments can be found in the figure legends including the statistical tests used, value of n, what n represents, number of experiments, and definition of center with precision measures.

