## Supplemental Information for "Developmental maturation of the hematopoietic system controlled by a Lin28b-*let*-*7*-PRC1 axis"

#### SUPPLEMENTAL FIGURE LEGENDS

Figure S1. Developmental maturation of the hematopoietic system.

(A) Frequencies of various HSC and MPP populations as a fraction of viable Lineage- c-kit+ Sca-1+ (LSK) cells in each hematopoietic organ. Data are presented as mean ± SEM with p values shown. Results aggregated over three independent experimental cohorts.

(B) Representative flow cytometry plots showing donor-derived engraftment at 4 weeks following transplantation of mononuclear cells from E14.5 FL, neonatal BM, or adult BM into congenic recipients.

(C) Quantification of lineage output from the indicated sources at the indicated times. P = not significant for all comparisons.

Figure S2. Developmental regulation of Lin28b and downstream targets in hematopoiesis.

(A) Heatmap showing expression of LIN28B and downstream targets in the indicated human HSC populations (Cesana et al., 2018).

(B) Heatmaps showing expression of Lin28b and downstream targets in Flk2- FL murine HSCs compared to adult bone marrow HSCs (Beaudin et al., 2016).

(C) Heatmaps showing expression of Lin28b and downstream targets in Flk2+ FL murine HSCs compared to adult bone marrow HSCs (Beaudin et al., 2016).

(D) Heatmaps showing expression of Lin28b and downstream targets in FL murine HSCs compared to adult bone marrow HSCs (Chen et al., 2019; Tober et al., 2018).

(E) ChIP-seq for H3K27ac in human FL or BM CD34+ HSCs/progenitors at the LIN28B locus (Huang et al., 2016).

(F) ChIP-seq for H3K27ac in human FL or BM CD34+ HSCs/progenitors at the CBX2 locus (Huang et al., 2016).

Figure S3. HSC and MPP populations in *Cbx2*$^{-/-}$ and control mice.

(A) Representative flow cytometry plots of E14.5 FL from the indicated genotypes.

(B) Quantification of HSC and MPP populations from the indicated genotypes. Data are presented as mean ± SEM with p values shown.

Figure S4. B-cell maturation in *Cbx2*$^{-/-}$ neonatal spleens.

(A) Representative flow cytometry plots of the indicated B-cell markers. Adult B-2 cells are CD5- B220-hi.

(B) Quantification of adult B-2 cells in the indicated genotypes. Data are presented as mean ± SEM with p values shown.

Figure S5. Analysis of PRC1 in HSPCs.

(A) CUT&RUN for the indicated histone modifications was performed in HUDEP-1 cells, and positions of mapped reads indicated.

(B) CUT&RUN analysis of HUDEP-1 cells for the indicated markers compared to ChIP-seq data from adult and fetal human CD34+ HSPCs at the indicated loci.

(C) CUT&RUN for the indicated histone modifications at the *Hoxa* gene cluster.

(D) CUT&RUN for the indicated histone modifications at the *Hoxb* gene cluster.

(E) CUT&RUN and ChIP-seq tracks for the indicated markers at the human *ERG* gene in either K562 cells or adult or fetal HSPCs. Candidate developmentally regulated enhancers are indicated.

(F) Representative transcription factor binding motifs with consensus sequences (top) compared to predicted motifs (bottom) in the enhancer sequences associated with the human *ERG* gene.

(G) H3K27Ac at the murine *Erg* enhancer (boxed) in adult and fetal mouse LSK cells(Chen et al., 2019).

Figure S1

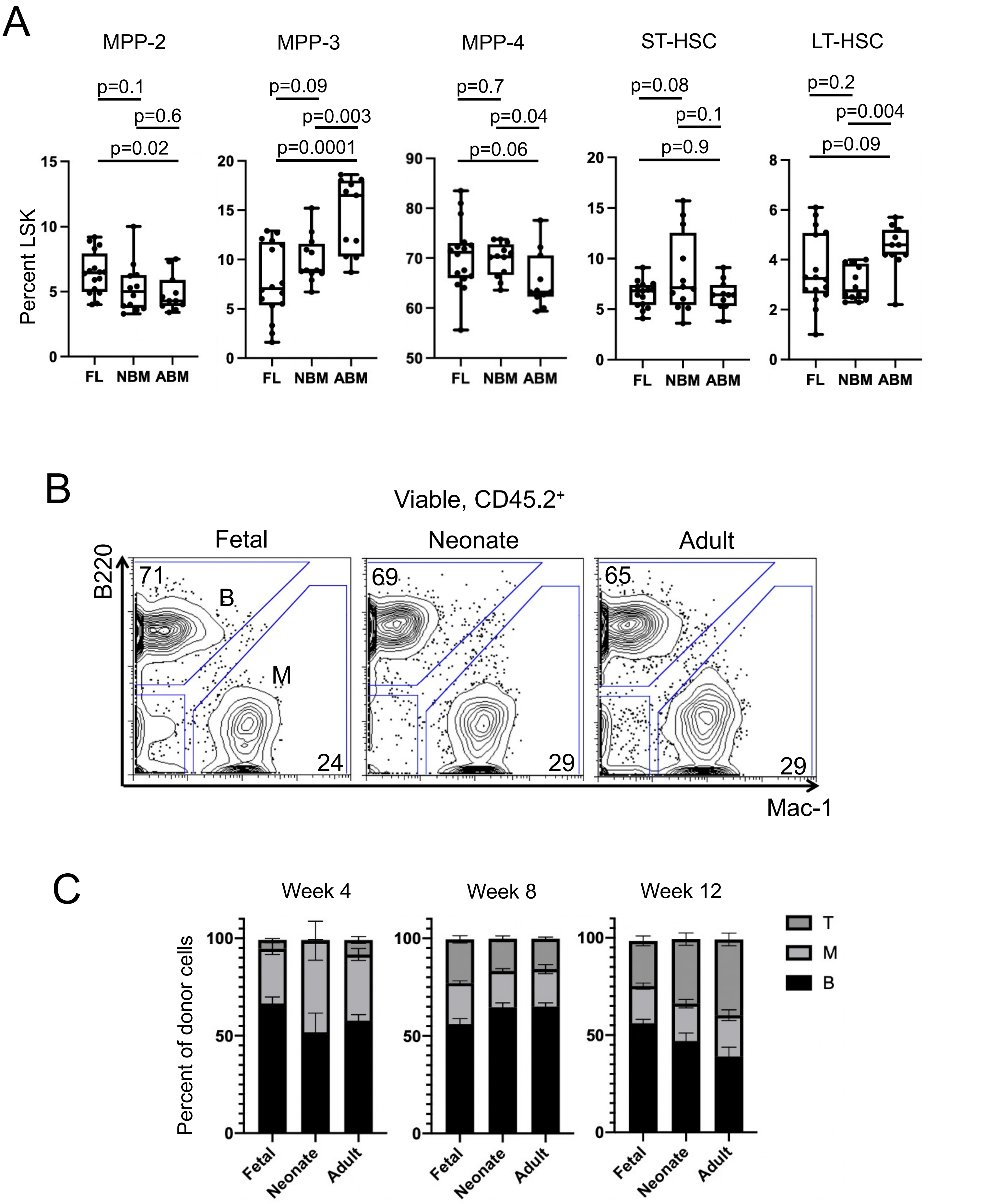
A
MPP-2
MPP-3
MPP-4
ST-HSC
LT-HSC
p=0.1
p=0.6
p=0.02
p=0.09
p=0.003
p=0.0001
p=0.7
p=0.04
p=0.06
p=0.08
p=0.1
p=0.9
p=0.2
p=0.004
p=0.09
Percent LSK
FL
NBM
ABM
B
Viable, CD45.2+
Fetal
Neonate
Adult
B220
Mac-1
71
24
69
29
65
29
B
M
C
Week 4
Week 8
Week 12
Percent of donor cells
Fetal
Neonate
Adult
T
M
B

Figure S2

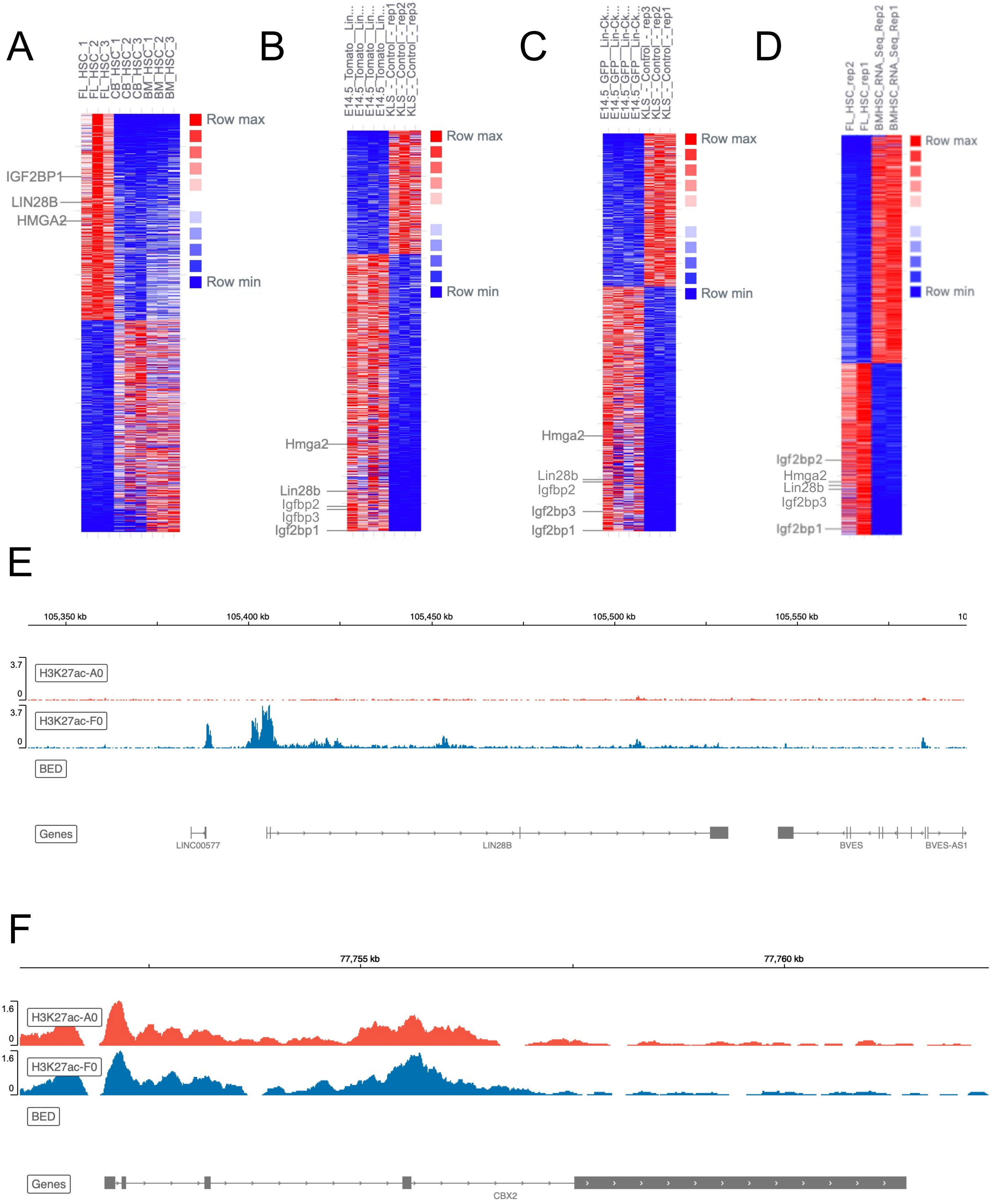
A
FL_HSC_1
FL_HSC_2
FL_HSC_3
CB_HSC_1
CB_HSC_2
CB_HSC_3
BM_HSC_1
BM_HSC_2
BM_HSC_3
Row max
Row min
IGF2BP1
LIN28B
HMGA2
B
E14.5_Tomato__Lin...
KLS_-_Control_-_rep1
KLS_-_Control_-_rep2
KLS_-_Control_-_rep3
Row max
Row min
Hmga2
Lin28b
Igfbp2
Igfbp3
Igf2bp1
C
E14.5_GFP__Lin-Ck...
KLS_-_Control_-_rep3
KLS_-_Control_-_rep2
KLS_-_Control_-_rep1
Row max
Row min
Hmga2
Lin28b
Igfbp2
Igf2bp3
Igf2bp1
D
FL_HSC_rep2
FL_HSC_rep1
BMHSC_RNA_Seq_Rep2
BMHSC_RNA_Seq_Rep1
Row max
Row min
Igf2bp2
Hmga2
Lin28b
Igf2bp3
Igf2bp1
E
105,350 kb
105,400 kb
105,450 kb
105,500 kb
105,550 kb
3.7
0
H3K27ac-A0
H3K27ac-F0
BED
Genes
LINC00577
LIN28B
BVES
BVES-AS1
F
77,755 kb
77,760 kb
1.6
0
H3K27ac-A0
H3K27ac-F0
BED
Genes
CBX2

Figure S3

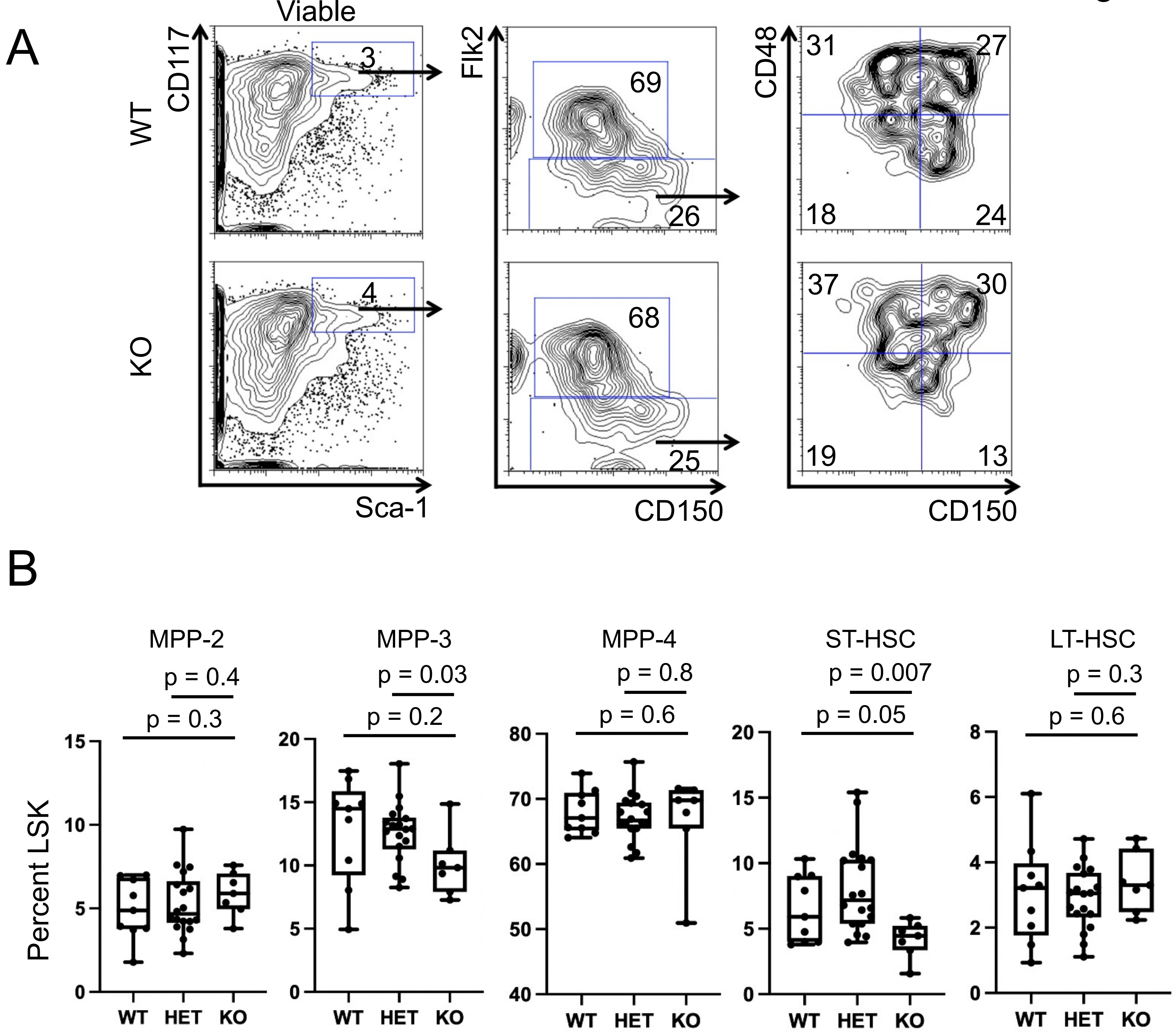
A
Viable
WT
KO
CD117
Sca-1
3
4
Flk2
CD150
69
26
68
25
CD48
31
27
18
24
37
30
19
13
B
Percent LSK
MPP-2
p = 0.4
p = 0.3
MPP-3
p = 0.03
p = 0.2
MPP-4
p = 0.8
p = 0.6
ST-HSC
p = 0.007
p = 0.05
LT-HSC
p = 0.3
p = 0.6
WT
HET
KO

Figure S4

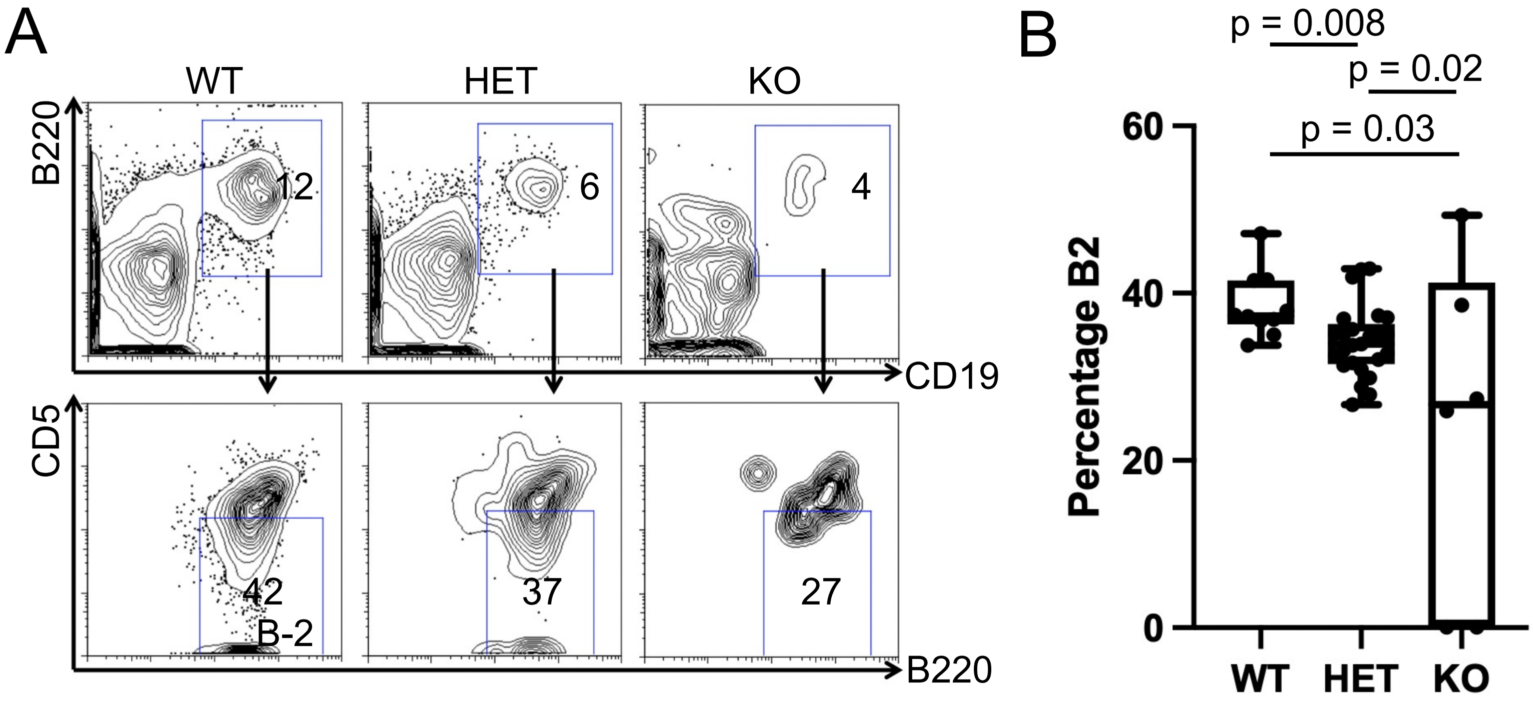
A
WT
HET
KO
B220
CD19
12
6
4
CD5
B220
42
B-2
37
27
B
p = 0.008
p = 0.02
p = 0.03
Percentage B2
60
40
20
0
WT
HET
KO

### Figure S5

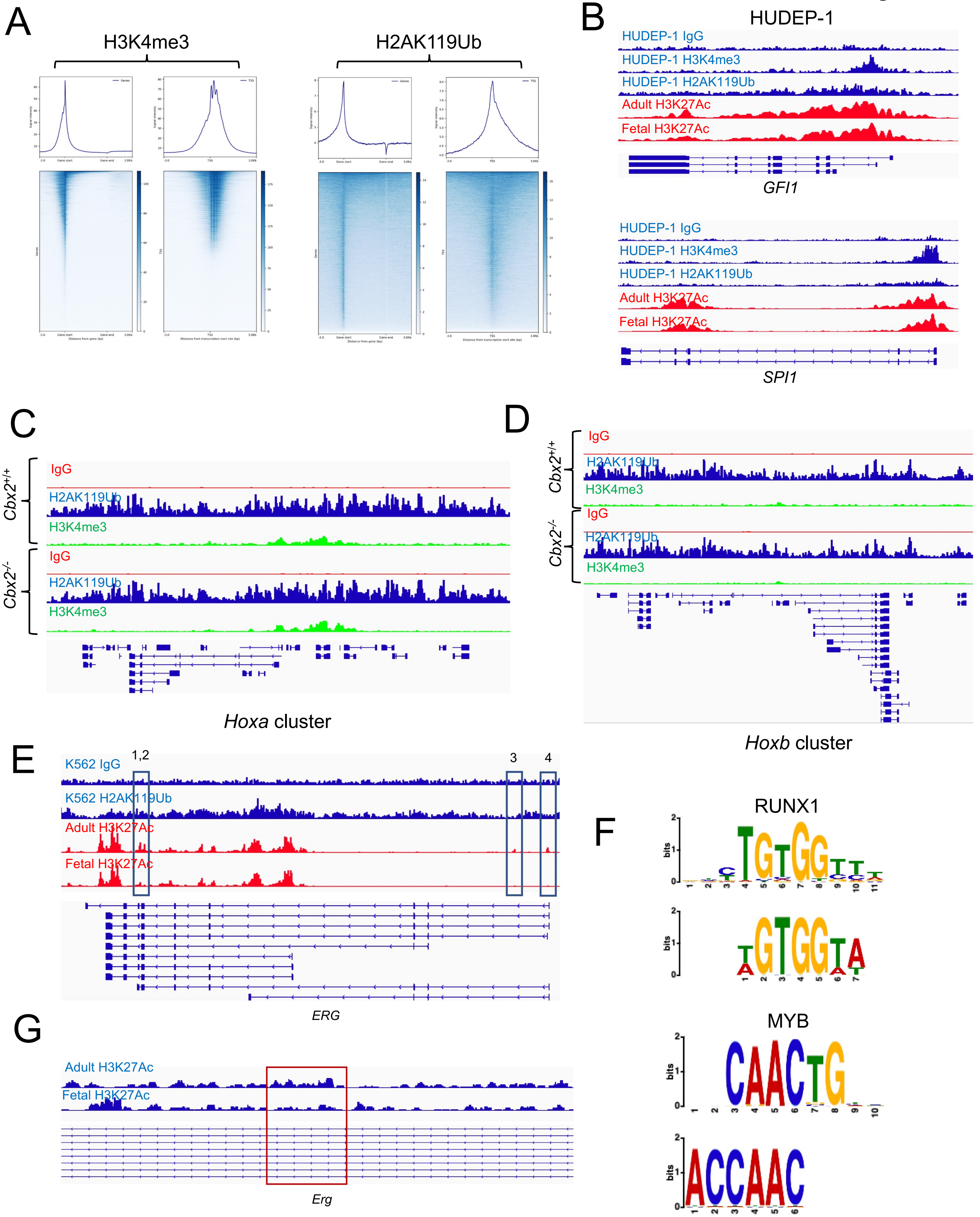
A
H3K4me3
H2AK119Ub
B
HUDEP-1
HUDEP-1 IgG
HUDEP-1 H3K4me3
HUDEP-1 H2AK119Ub
Adult H3K27Ac
Fetal H3K27Ac
GFI1
HUDEP-1 IgG
HUDEP-1 H3K4me3
HUDEP-1 H2AK119Ub
Adult H3K27Ac
Fetal H3K27Ac
SPI1
C
$Cbx2^{+/+}$
IgG
H2AK119Ub
H3K4me3
$Cbx2^{-/-}$
IgG
H2AK119Ub
H3K4me3
Hoxa cluster
D
$Cbx2^{+/+}$
IgG
H2AK119Ub
H3K4me3
$Cbx2^{-/-}$
IgG
H2AK119Ub
H3K4me3
Hoxb cluster
E
1,2
3
4
K562 IgG
K562 H2AK119Ub
Adult H3K27Ac
Fetal H3K27Ac
ERG
F
RUNX1
MYB
bits
G
Adult H3K27Ac
Fetal H3K27Ac
Erg
